# Small interfering RNAs are highly effective inhibitors regarding Crimean-Congo hemorrhagic fever virus replication *in vitro*

**DOI:** 10.1101/2020.05.13.093047

**Authors:** Fanni Földes, Mónika Madai, Henrietta Papp, Gábor Kemenesi, Brigitta Zana, Lili Geiger, Katalin Gombos, Balázs Somogyi, Ildikó Bock-Marquette, Ferenc Jakab

**Author notes:** Correspondence (F.J.); /29044 (F.J.).

## Abstract

Crimean-Congo hemorrhagic fever virus (CCHFV) is one of the prioritized diseases of World Health Organization, considering its potential to create a public health emergency and more importantly, the absence of efficacious drugs and/or vaccines regarding treatment. The highly lethal nature characteristic to CCHFV restricts research to BSL-4 laboratories, which complicates effective research and developmental strategies. In consideration of antiviral therapies, RNA interference can be used to suppress viral replication by targeting viral genes. RNA interference uses small interfering RNAs (siRNAs) to silence genes. The aim of our study was to design siRNAs that inhibit CCHFV replication and can serve as a basis for further antiviral therapies. A549 cells were infected with CCHFV after transfection with the siRNAs. Following 72 hours, nucleic acid from the supernatant was extracted for Droplet Digital PCR analysis. Among the investigated siRNAs we identified four effective candidates against all three segments of CCHF genome: one for the S and M segments, whilst two for the L segment. Consequently, blocking any segment of CCHFV leads to changes in the virus copy number that indicates an antiviral effect of the siRNAs *in vitro*. The most active siRNAs were demonstrated a specific inhibitory effect against CCHFV in a dose-dependent manner. In summary, we demonstrated the ability of specific siRNAs to inhibit CCHFV replication *in vitro*. This promising result can be used in future anti-CCHFV therapy developments.

## 1. Introduction

Crimean-Congo hemorrhagic fever virus (CCHFV) categorically belongs to the *Orthonairovirus* genus, the *Nairoviridae* family in the *Bunyavirales* order. CCHFV is causing a mild to severe hemorrhagic disease in humans, with fatality rates from 5% up to 30% [1].

CCHFV is characterized by a tripartite single-stranded RNA genome (S, M and L segment) of ambisense (S) and negative (M, L) polarity. The three genome segments encode four structural proteins: the RNA dependent RNA polymerase is encoded by the large (L) segment, the glycoproteins (G_N_ and G_C_) are encoded by the medium (M) segment, and the nucleocapsid protein and nonstructural protein are encoded by the small (S) segment [2].

Emerging infectious diseases (EIDs) are growing threats to animal and human health. CCHFV is a tick-borne pathogen that causes an increasing number of severe infections and presents over a wide geographic range, including areas in South-Eastern Europe, Western and Central Asia, the Middle East and Africa as well [1]. This virus is transmitted primarily by ticks, but the spectrum of natural hosts for CCHFV includes a wide variety of domestic and wild animals [3].

There are neither vaccines nor effective antiviral therapies for the treatment of CCHFV infections in humans to date [4]. There is a growing need for advantaged research and development activities for such pathogens as CCHFV, since there is a constantly growing geographic and epidemiologic burden of the disease and BSL-4 capacity is limited throughout the world, which can safely handle such research.

Among antiviral therapies, RNA interference (RNAi) can be used to suppress viral replication by targeting either viral- or host genes that are needed for viral replication. Since its discovery in 1998 [5], it has revolutionized the mechanism of gene silencing and improved our understanding of the endogenous mechanism of gene regulation to enhance the use of new tools for antiviral research. Silencing viral genes such as viral polymerases, master regulators of viral gene transcription and viral genes that act early in the viral life cycle, may suppress viral replication more effectively than targeting late or accessory viral genes. Moreover, RNAi could target viral proteins and pathways, which are unique to the viral life cycle and it has become possible to interfere with viral infections and replication without unacceptable host cell toxicity [6]. Accordingly, the major advantage of RNA interference is its target specificity. In recent years, many viruses have been successfully targeted by RNA interference such as human immunodeficiency virus (HIV) [7,8], Severe Acute Respiratory Syndrome coronavirus (SARS-CoV) [9], Hepatitis B virus (HBV) [10], Hepatitis C virus (HCV) [11], Influenza A virus [12], Hazara virus (HAZV) [13], Langat virus (LGTV) [14], Andes virus (ANDV) [15] and West Nile virus (WNV) [16]. So far, to the best of our knowledge, this is the first study that used RNA interference to inhibit CCHFV replication *in vitro*. Although, another member of the *Nairoviridae* family (e.g. HAZV) was already researched on virus gene silencing by RNA interference [13].

RNA interference uses small double-stranded RNAs with a complementary sequence to the target silencing genes. Nevertheless, endogenous gene silencing operates through multiple mechanisms such as mRNA cleavage, inhibition of translation, and epigenetic modifications of chromatin, of which mRNA cleavage is the most efficient mechanism for antiviral therapies [6]. Small interfering RNAs (siRNA) are the active agents in RNA interference. The siRNAs are 21–22 nucleotides long, serve as a guide for cognate mRNA degradation [17]. Naturally, these siRNAs are a result of endonucleolytic processing of a larger precursor RNA. Experimentally, RNAi can be triggered in mammalian cells after the transfection of synthetic siRNA using suitable transfection reagents. These siRNAs are incorporated into a cytoplasmic RNA-induced silencing complex (RISC) which cleaves exogenous double-strand siRNAs and leaving an unpaired guide strand to search for complementary mRNAs. If the target site on the mRNA has nearly perfect complementarity to the guide siRNA, the mRNA is cut by an Argonaute (Ago) endonuclease in the RISC and is degraded. This way, siRNA is silencing the expression of the protein encoded by the target mRNA. Typically, protein expression is reduced but not eliminated [6].

Recent works have shown that more effective antiviral therapies are urgently needed to treat virus infections especially for viruses with growing epidemic potential [9,15]. Furthermore, these RNA interference experiments have shown that the application of siRNAs can inhibit viral infection by targeting viral genes [18,19]. However, many aspects of the CCHFV cell entry, replication and pathogenesis remain poorly defined. It was mostly studied by using minigenome systems or virus-like-particle systems considering its highly infectious nature and the lack of BSL-4 laboratories [2]. In our present study, we aimed to design chemically synthesized siRNAs that can inhibit first time CCHFV replication *in vitro*. This study presents the first step forward to future RNAi-based CCHF antiviral therapy development.

## 2. Materials and Methods

### 2.1. Cell line, virus amplification and titer determination

A549 cells (human lung carcinoma cell line, ATCC CCL-185) were grown in Dulbecco’s modified eagle medium (DMEM) (Lonza) supplemented with 10% heat-inactivated fetal bovine serum (FBS) (EuroClone) and 1% Penicillin-Streptomycin (Lonza) maintained at 37°C in a humidified atmosphere containing in a 5% CO_2_.

A549 cells with 60% confluence were infected by the CCHFV Kosova Hoti strain [20] in our experiments. The virus was grown to high titers on A549 cells and the supernatants were aliquoted and were frozen at −80°C in 1 ml vials and constituted the viral stock. All laboratory manipulations associated with infectious CCHFV were performed in a BSL-4 suite laboratory, aligned to the University of Pécs, Szentágothai Research Centre.

CCHFV viral stock was titrated using the TCID50 method with the immunofluorescence assay. Briefly, serial 10-fold dilutions of CCHFV supernatant were inoculated (100 µl) on 60% confluent A549 cells (30000 cells/well) in 48-well plates. Viral adsorption was allowed for 1 hour at 37°C. After washing cells with PBS three times, cells were incubated for 3 days at 37°C in DMEM supplemented with 2% FBS. The fixation and the immunofluorescence assay were performed as previously described using with polyclonal mouse antibody which was produced against the recombinant CCHFV capsid protein [21]. The percentage of infected cells was observed with immunofluorescence microscopy and recorded for each virus dilution then results were used to mathematically calculate a TCID50 result with the Spearman-Karber method. During our experiments, A549 cells were infected with CCHFV at a MOI of 0.1 in our following infection and transfection assays.

### 2.2. Design and synthesis of siRNAs

The sequences of CCHFV Kosova Hoti strain S, M and L genomic segments (GenBank: DQ133507, EU037902, EU044832) were used to design the siRNAs. Synthetic 21-nucleotide siRNAs with short 3’ overhangs (UU) were designed by the Whitehead siRNA Selection Program to have an antisense strand complementary to the CCHFV [22]. The siRNA sequences were chosen according to the algorithm score. For each viral mRNAs, five siRNAs were synthesized by Dhramacon™ (Table 1). Sequences were subjected to a BLAST search against GenBank to minimize off-target effects. All lyophilized siRNAs were reconstituted according to the manufacturer’s instruction, aliquoted in 10 µM stock solutions and were stored at −20°C until further use. The TOX siRNA (siTOX) (Dharmacon™ RNAi technologies, Lafayette, USA) was used to determine transfection efficacy.

**Table 1.**
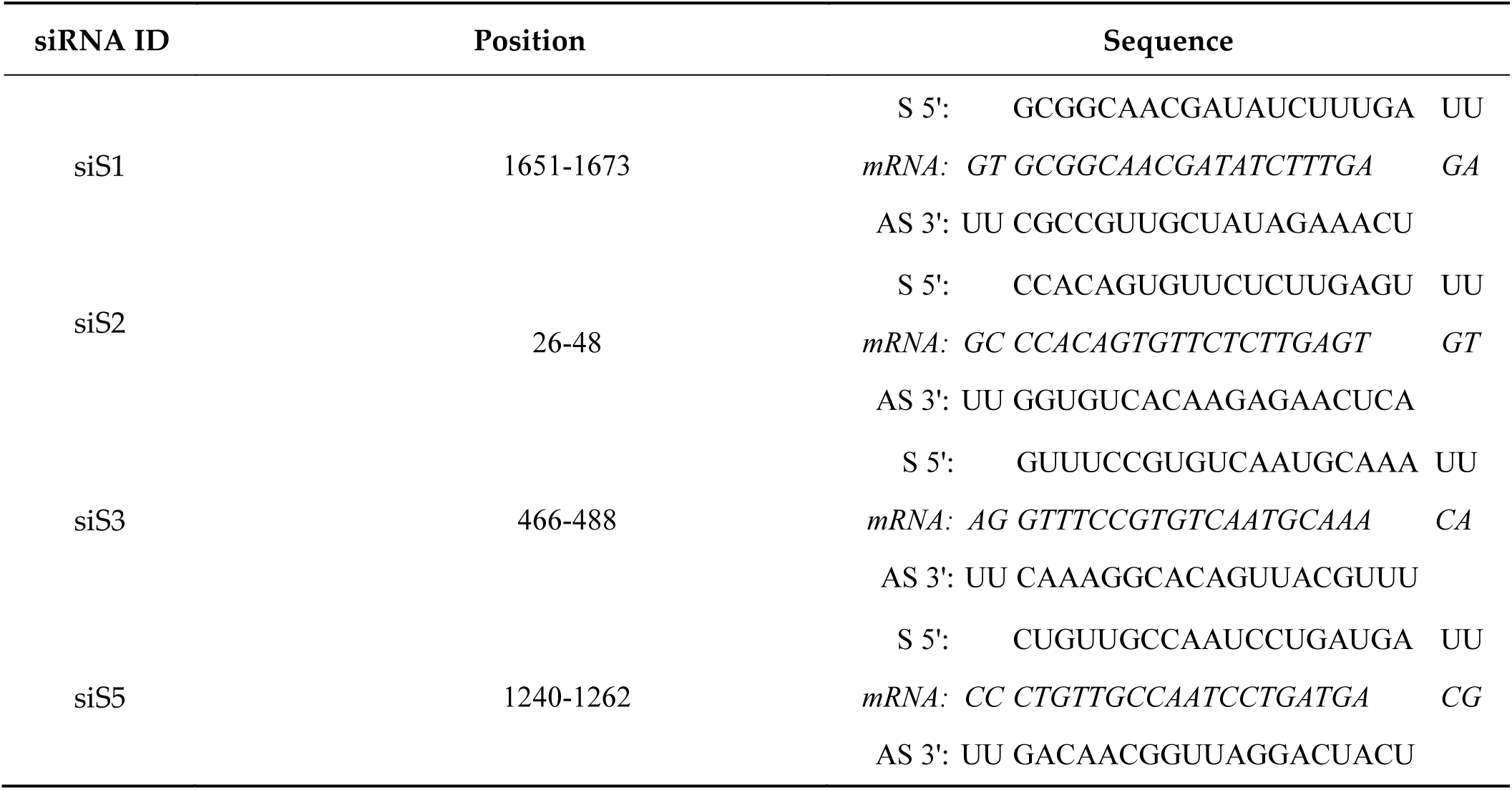
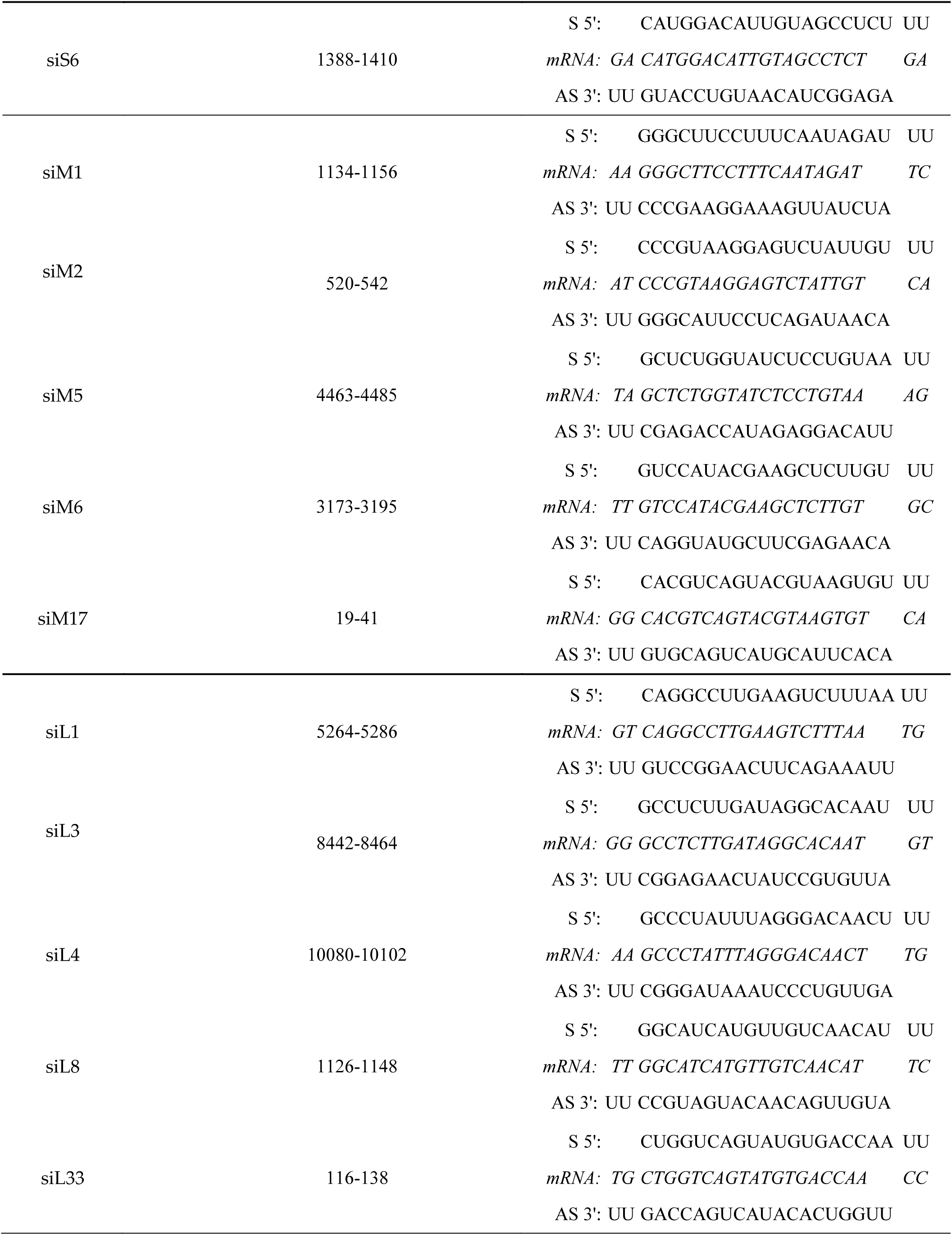
List of designed siRNAs

### 2.3. Transfection efficiency

For each experiment, transfection efficiency was monitored by transfecting A549 cells with 200 nM of siTOX (Dharmacon™) under the same experimental conditions. Cells successfully transfected with siTOX went under apoptosis and cell death within 24-48 hours. After 3 days of incubation, siTOX transfected cells were trypsinized and manually counted using a hematocytometer (Trypan blue exclusion assay). Transfection efficiency was calculated as the ratio between the numbers of viable siTOX-transfected cells versus non-transfected cells. In our experiments, we experienced an average of 80% transfection efficiency.

### 2.4. Cytotoxicity tests

In some cases, the designed siRNAs could interfere with the tested cells’ genes (off-target effect) and cause cell death. During the concentration-dependent transfection, microscopic observation was performed. A549 cells were transfected with different concentrations (ranging from 0.1 nM to 300 nM) of siRNAs. Cells were observed microscopically after the transfection at 24, 48 and 72 hours. During the trypan blue exclusion assay, cell deaths and cell morphological changes have been recorded if the siRNAs targeted S, M or L segments of CCHFV at high siRNA concentration.

Besides microscopic observation, cell cytotoxicity was examined with a luminescence cell viability assay kit (Promega – Cell Titer Glo Luminescent assay). This method determines the number of viable cells in culture, based on quantitation of the ATP present. Cells were transfected with different concentrations of siRNAs (ranging from 0.1 nM to 200 nM). After 72 hours of transfection, luminescence measurement was performed. The IC50 was calculated using GraphPadPrism version 8.00 software (Graph Pad Software, San Diego California, USA) for non-linear regression.

The use of cytotoxicity tests was important to find out the concentration at which siRNAs do not cause cell death but their concentration is high enough to inhibit virus replication.

### 2.5. Transfection and infection assay

Transfection and infection experiments were performed on A549 cells in the BSL-4 laboratory. A549 cells were seeded in 96-well plates at a density of 2×10^4^ cells/well to achieve 60-70% confluent cell monolayers on the day after in a humidified incubator at 37°C with 5% CO_2_.

Cells were transfected in triplicate biological replicates with siRNAs in the following final concentrations: 10 nM, 50 nM, 200 nM. Various siRNA concentrations were complexed with the transfection reagent Lipofectamine RNAiMax transfection reagent (Thermo Fisher Scientific) according to the manufacturer’s instructions. The transfection reagent and siRNAs were diluted in Opti-MEM medium (Gibco). The final volume of Lipofectamine RNAiMax was 1.5 µl/well. The transfection mixture was incubated for 20 minutes at RT to allow the formation of siRNA-lipid complexes and 100 µl of the solution were added slowly dropwise to each well. Mock transfected, non-transfected A549 cells were used as controls for the experiments. Mock-transfected cells go through the transfection process without the addition of siRNA while non-transfected cells have not been treated at all. 16 hours post-transfection, cells were gently washed twice with DMEM. Thereafter transfected cells were infected with CCHFV at a MOI of 0.1. The inoculum was incubated for 1 hour to allow the absorbing of the virus on transfected cells. Cells were then cultivated in DMEM supplemented with 2% FBS, 1% Penicillin-Streptomycin for 48 hours. Non-transfected A549 cells which were infected with CCHFV at a MOI of 0.1 were used as positive cell controls. Cell morphology was monitored and 200 µl cell supernatant was harvested before nucleic acid extraction.

Virus replication decrease was assessed by determining the number of genome copies in 200 µl cell supernatant by qRT-PCR and RT-ddPCR.

### 2.6. Viral RNA extraction and qRT-PCR for pre-screening

To investigate the inhibitory effect of all designed siRNAs in different concentrations (ranging from 10 nM to 200 nM), firstly qRT-PCR assay was performed as a pre-screen.

Template viral RNA from transfected cells and control cells were extracted from 200 µl culture supernatant using DNA/RNA extraction kit (Geneaid), according to the manufacturer’s protocol. The nucleic acid extraction was performed in the BSL-4 suite laboratory. The RNA elution was done in a volume of 50 µl of elution buffer and was stored at −80°C until further use.

The quantitative real-time TaqMan based assay was carried out using a One-step RT-PCR kit (Qiagen) in the Light Cycler 2.0 system (Roche). CCHFV specific primers and probe were based on Atkinson et al. publication (Table 2)[23]. Reaction profile was as follows: reverse transcriptions at 50°C for 30 minutes, initial denaturation at 95°C 15 minutes, followed by 50 cycles of amplification at 94°C 15 seconds, 51°C 30 seconds and 72°C 20 seconds.

**Table 2:**
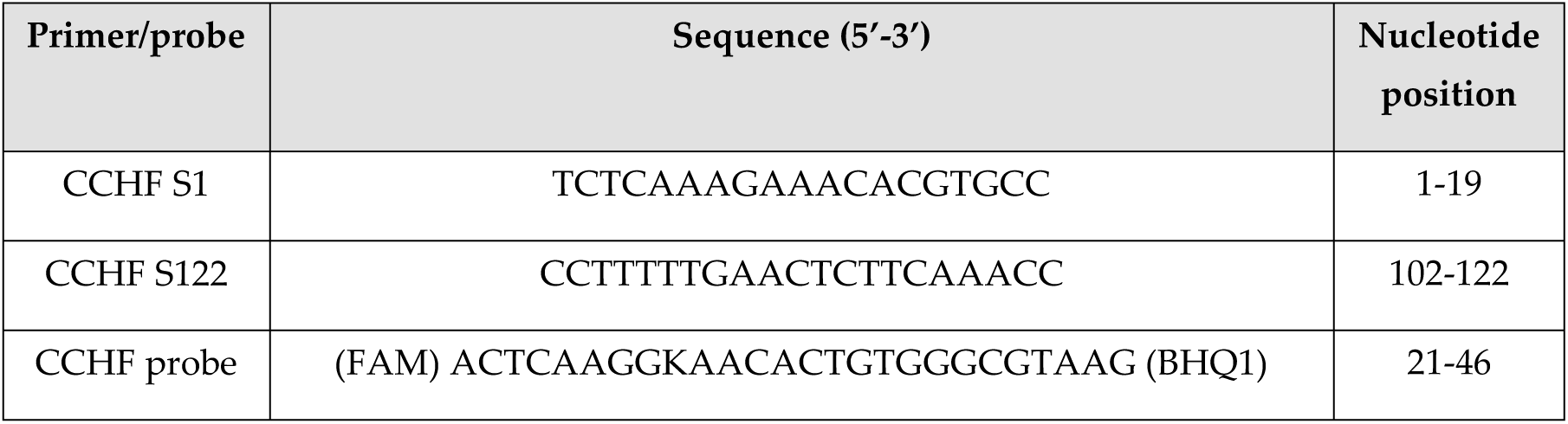
Primers and probe information for the CCHF real-time RT-PCR assay based on Atkinson et al. publication

### 2.7. Droplet digital RT-PCR and data analysis

After RT-PCR prescreening, the siRNAs which inhibited CCHFV replication effectively were measured by RT-ddPCR in three time biological repetitions with different concentrations (ranging from 10 nM to 200 nM).

QX200 Droplet Digital PCR system (Bio-Rad, CA, USA) was used to determine CCHFV copy number decrease triggered by siRNAs from supernatants. One-Step RT-ddPCR advanced kit for probes (Bio-Rad, CA, USA) was used in our experiments. The RT-ddPCR reaction mixture consisted of 5 µl of a ddPCR Supermix, 2 µl reverse transcriptase, 1 µl 300 mM DTT, 900 nM CCHFV specific primers and 250 nM probe, 1 µl of sample nucleic acid solution and nuclease-free H_2_O in a final volume of 22 µl. The final concentrations of CCHFV specific primers and probe [23] were the same as for RT-qPCR assays. The entire reaction mixture was loaded into a disposable plastic cartridge (Bio-Rad, CA, USA) together with 70 µl of droplet generation oil for probes (Bio-Rad, CA, USA) and placed in the QX200 Droplet Generator (Bio-Rad, CA, USA). After processing, the droplets generated from each sample were transferred to a 96-well PCR plate (Bio-Rad CA, USA) and heat-sealed with PX1TM PCR Plate Sealer (Bio-Rad, CA, USA). PCR amplification was carried out on a C1000 TouchTM Thermal Cycler with 96-Deep Well Reaction Module (Bio-Rad, CA, USA) using a thermal profile of beginning at reverse transcription: 50°C for 1 hour and 95°C for 10 min, followed by 40 cycles of 95°C for 30 s and 55°C for 60 s, 1 cycle of 98°C for 10 min, and ending at 4°C. After amplification, the plate was loaded on the QX200 Droplet Reader (Bio-Rad, CA, USA) and the droplets from each well of the plate were read automatically. Positive droplets, containing amplification products, were partitioned from negative droplets by applying a fluorescence amplitude threshold in QuantaSoftTM analysis software (Bio-Rad, CA, USA). The threshold line was set manually at 3780 amplitudes for every sample. Quantification of the target molecule was presented as the number of copies per µl of the PCR mix. All siRNAs in different concentrations were tested in three biological replicates. During the PCR reactions (qPCR and ddPCR) the same target segment was used.

### 2.8. Statistical analysis

All experiments were repeated in three biological replicates. In our study, we compared the antiviral effect of selected effective siRNAs in different concentrations to the positive control to detect significant variations using the Student’s t-test. The measured dataset was statistically analyzed in the R environment [24]. The bar plots were created with ggplot2 R package [25]. During PCR reactions (qPCR and ddPCR), three biological replicates of siRNA inhibited CCHFV samples were used and we did not use technical replicates in case of these siRNAs inhibited CCHFV samples since the three biological replicates include the technical replicate. However, the controls were used in three biological and three technical repeats.

## 3. Results

In our study, 15 siRNAs were designed and synthesized to test the inhibitory activity on CCHFV replication and target the mRNAs produced by S, M and L segments. We analyzed the high inhibitory effect of some S (siS2), M (siM1) and L (siL3, siL4) segment-specific siRNAs. We experienced that siRNAs inhibited CCHFV replication in different efficiency and dose-dependent manner.

### 3.1. Cytotoxicity tests

During the experiments, two different types of cell viability tests were used: light microscopic observation and luminescence cytotoxicity measurement (Table 3).

The siRNAs treatment could cause visual cytopathogenic effects (CPEs) and affect viral growth, therefore we performed light microscopic observation to evaluate cell growth and viability. Firstly, we had to find the appropriate siRNA concentration that is effective in inhibiting CCHFV replication but not toxic to the cells. In our cytotoxicity experiments, after three days of siRNAs transfection, the cell number per well was observed and compared to non-transfected cells by manual counting with hematocytometer. In these experiments, we did not detect the cytotoxic effect of siRNAs on A549 cells at any lower concentrations used. However, a high concentration of siRNAs (300 nM) caused cell morphology changes and cell death.

Besides morphological observation with light microscopy, luminescence cytotoxicity measurements (Promega – Cell Titer Glo Luminescent assay) were used. The half-maximal inhibitory concentration (IC50) is used to measure the potency of a material inhibiting a specific biological function. IC50 is a quantitative measurement that indicates how much of a particular inhibitory substance (e.g. siRNA) is needed to inhibit given biological process by 50% *in vitro*. The IC50 was calculated using GraphPadPrism version 8.00 software (Graph Pad Software, San Diego California, USA) for non-linear regression. In most cases, during cell viability tests, results that were observed microscopically were the same as the luminescence cytotoxicity measurements. Cell control and transfection reagent control showed the luminescence cytotoxicity measurements indicating no cell cytotoxicity of transfection reagent. In the case of siRNAs against the CCHFV S segment, the results were the same with the cell viability tests: IC50 was observed around 200 nM in every siRNAs against the S segment. The most efficient S segment siRNA (siS2) IC50 value was 246.7 nM which was calculating by GrapPhadPrism version 8.00 software. CCHFV M segment siRNAs were proven to be non-toxic for the A549 cells up to 250 nM concentration. The most efficient M segment siRNA (siM1) IC50 value was 251.8 nM. However, siRNAs against the M segment were used in 200 nM concentration because of the comparability. In the case of siRNAs against the CCHFV L segment results were the same with the cell viability tests, therefore we used them at maximum 200 nM concentration in our further experiments. The most efficient L segment siRNAs (siL3, siL4) IC50 values were 180.92 and 180.8 nM, respectively.

Summarizing the cell viability results, the minimum concentration of siRNAs was set to 10 nM and the maximum concentration to 200 nM (10 nM, 50 nM, 100 nM, and 200 nM) in case of every segment. The little differences found between the cytotoxicity tests indicate that the use of microscopic observation alone is not sufficient enough to detect cell viability and specify the appropriate concentration of siRNAs.

### 3.2. Inhibition of CCHFV replication using segment-specific siRNAs

qRT-PCR TaqMan assay was performed as pre-screening because of the large sample size, cost and time effectiveness. The siRNAs which showed promising inhibitory effect against CCHFV based on qRT-PCR results were chosen for further experimentation. Out of the 15 siRNAs that were designed against CCHFV, 9 were selected. In our further RNA interference experiments, three siRNAs for every segment of CCHFV (siS1, siS2, siS6, siM1, siM6, siM17, siL1, siL3, and siL4) were used (Figure 1/A).

**Figure 1.**
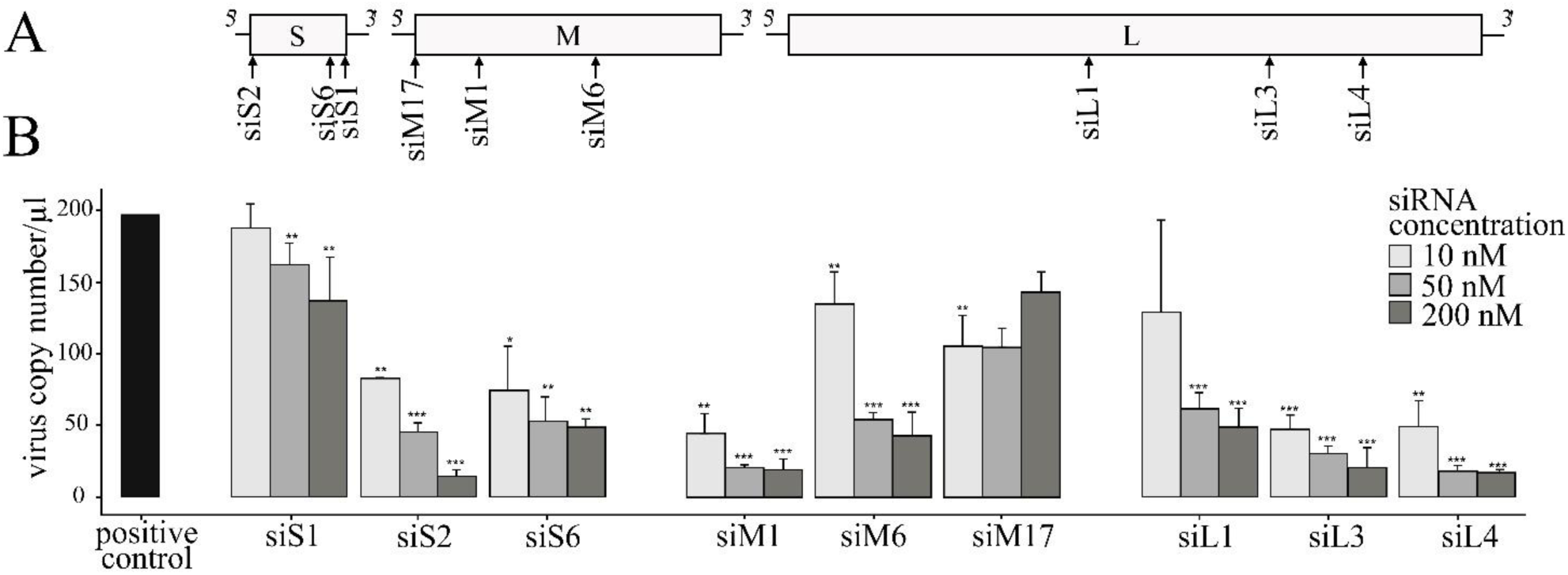
A549 cells were transfected with siRNAs which were designed for CCHFV S, M and L segments in different concentrations (10 nM, 50 nM, and 200 nM). After transfection, cells were infected with CCHFV at a MOI of 0.1. Three biological replicates of siRNA inhibited CCHFV samples were used and the positive control was used in three biological and three technical repeats. The virus copy number was determined after 72 hours by RT-ddPCR. (a) CCHFV schematic gene map is containing designed CCHFV-specific siRNAs site; (**b**) Inhibitory effect of siRNAs against CCHFV: the horizontal axis represents the virus copy number/µl and the vertical axis represents the positive control and designed siRNAs. Student’s t-tests were significant if *: *P* < 0.05, **: *P* < 0.01, ***: *P* < 0.001. Error bars represent the standard deviation (SD) of the means for three independent experiments.

Based on the ddPCR results, a high and significant copy number decrease in the case of some siRNAs (siS2, siM1, siL3 and siL4) was detected. As shown in Figure 1/B, when siS2 was used at 200 nM concentration, it has strongly and significantly inhibited CCHFV replication compared to the positive control (*P*<0.001). Among siRNAs against CCHFV S segment, siS2 was the most efficient inhibitory siRNA. Furthermore, siS6 has shown a moderate but significant inhibitory effect in CCHFV replication (*P*<0.01). In contrast, significant antiviral inhibitory effect of siS1 at 10 nM concentration was not detected. Between siRNAs which were designed for the M segment, siM1 had strong and significant antiviral activity at 100 nM concentration (*P*<0.001). Moreover, siM6 has also shown CCHFV inhibitory effect at medium level (*P*<0.001). In contrast, siM17 has not inhibited CCHFV replication significantly at 200 nM concentration. In case of the L segment, when siL4 was used at 200 nM concentration, it has strongly inhibited CCHFV replication compared to the positive control (*P*<0.001). SiL3 has also shown significant, high activities on CCHFV replication, while siL1 has shown moderate efficiency. (*P*<0.001) (Figure 1/B).

At least one highly inhibitory siRNA was found in case of every segment. The siRNAs that were designed for the S segment: the siS2 has shown the most efficient which decreased the virus copy number by about 93% at 200 nM concentration. In case of M segments siRNAs, siM1 has decreased the virus copy number by about 90% at 200 nM concentration which was almost the same as siS2. Among siRNAs that were designed to L segment, siL4 has affected CCHFV replication (decrease by about 92%) at 200 nM concentration just like siS2 and siM1. Inhibitory effect against CCHFV was not caused by siS1 and siM17.

QuantaSofts’ RT-ddPCR raw fluorescence readouts have shown negative and positive controls in Figure 2. A negative droplet population was shown by the negative control sample without any positive droplets. The positive control sample has appeared as a massive positive droplet population above the threshold level. In case of the positive control sample, the positive droplet “rain” was caused by the high concentration of CCHFV and appeared as a background signal. Concentration-dependent high inhibitory effect was shown by siS2 and siM1. At 10 nM concentration, positive droplet number was high in case of siS2, however, at 200 nM concentration, positive droplet number was decreased extensively. SiM1 acted similarly as siS2. A medium inhibitory effect against CCHFV replication was presented by SiM6. The positive droplet number decreased moderately from 10 nM to 200 nM compared to siS2 and siM1 events. In case of siM17, the significant inhibitory effect was not detected.

**Figure 2.**
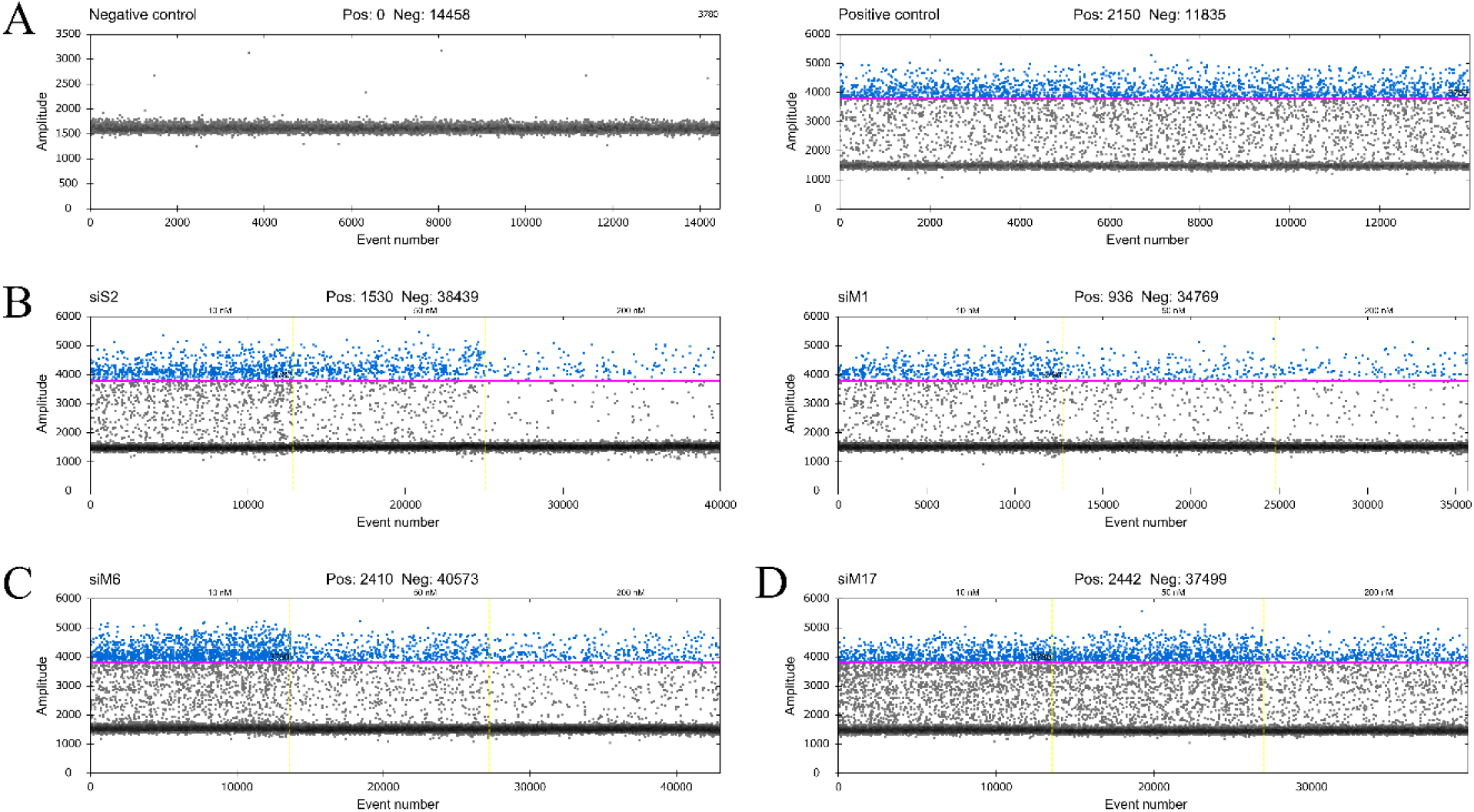
QuantaSofts’ RT-ddPCR fluorescent readouts. The horizontal axis represents the event number and the vertical axis represents the fluorescence amplitude in the FAM channel. The strict threshold line (pink line) was set for every sample at 3780 amplitude. Positive droplets were represented in blue and negative droplets were represented in grey. (A) Negative control sample was shown as the negative droplet population without positive droplets, positive control sample was shown as the extensive positive droplet population; (B) siS2 and siM1 respectively were shown a concentration-dependent high inhibitory effect, different concentrations (10 nM, 50 nM, 200 nM) were separated with yellow, dotted line; (C) siM6 has shown concentration-dependent medium inhibitory effect at different concentrations (10 nM, 50 nM, 200 nM) that were separated with yellow, dotted line; (D) siM17 has shown low inhibitory effect against CCHFV at different concentrations (10 nM, 50 nM, 200 nM) that were separated with yellow, dotted line.

## 4. Discussion

Therapeutic options for the treatment of CCHFV infection are lacking, with the noticeable exception of ribavirin, which is currently recommended by the WHO. Nevertheless, novel and more sophisticated antiviral therapies against nairovirus infections are urgently needed. In the last few years, several studies have shown that siRNAs have the potential to be operated as a specific therapeutic strategy against some viral infections [11,14,26,27]. However, most of these experiments are in the *in vitro* studies and translating RNAi in clinic, as a conventional treatment option remains a pivotal challenge. In case of *in vivo* therapies, one of the most difficult parts is efficiently and specifically delivering siRNA to target tissues and cells. Moreover, the poor cellular uptake of siRNAs in combination with rapid enzymatic degradation are limiting RNAi usage *in vivo* therapies. Besides, different classes of siRNA chemically modifications can increase the efficiency of delivery. Fortunately, despite difficulties in virus entry, cytotoxicity and the stimulation of unspecific immune response researches evolved and reached *in vivo* experiments [18].

We baselined our control strategy exclusively to the results of cytotoxicity tests without using non-targeted siRNA as seen in other related papers as well [28]. We evaluated the antiviral activity of siRNAs targeting the S (nucleoprotein), M (glycoproteins) and L (polymerase) transcripts of CCHFV for the first time *in vitro*. The siRNAs were designed for each CCHFV segment in an effort to find the most effective ones. We observed that among all tested siRNAs, almost half of them (siS2, siS6, siM1, siM6, siL1, siL3, siL4) were capable of reducing CCHFV copy number by more than about 70% during *in vitro* infection studies, comparing to the positive control. However, strong inhibition of CCHFV replication (by about 90%) was performed by only four siRNAs (siS2, siM1, siL3, and siL4). The unusual ability of many siRNAs inhibits the virus, contrary to previous studies, is due to the successful design and the high rate of transfection achieved.

In case of the CCHFV S segment protein, nucleoproteins play a central role in the regulation of viral replication. Nucleoprotein associated with genomic viral RNA to form RNPs and provided as a template for the polymerase. In the last few years, several homologous interferences have been described as the inhibition of S segment of other nairoviruses by siRNAs and suppression of viral replication [15,26,27]. Levin et al. found that Akabane virus (AKAV) infected Vero cells indicated more than 99% inhibition [26], whilst, Chiang et al. described siRNA against the S segment of andes virus (ANDV) greatly reduced levels (>60%) of viral protein expression [15]. In case of the hazara virus inhibition, the siRNAs which were designed against the S segment had a higher effect (up to 90%) than those targeting M and L segments [27]. Several experiments performed has shown that the S segment of genus *Orthobunyavirus* is the RNA interference prime target in arthropod cells [29,30]. In our study, among siRNAs that were designed against the S segment, siS2 has inhibited effectively (93%) CCHFV copy number. Our study is in agreement with previous works [15,27] that targeting the S segment by siRNAs can produce an effective inhibitory impact. In consequence, using the S segment as the target for silencing virus replication has proven to be an option for future therapeutics. Hereafter, using siRNAs together can have superior effect against virus infections [26]. Our plans include the combined use of designed siRNAs against CCHFV infection.

CCHFV glycoproteins (Gn, Gc) are involved in cell entry, initial binding and fusion. However, the details of specific glycoprotein involvement remain unknown [2]. In contrast with other studies, a high inhibitory effect of siRNA (90%) was found against the M segment. Furthermore, Chiang et al. described that viral glycoproteins are limiting factors for virus production and viral glycoproteins are detected mainly in the lysosome rather than on the cell surface in genus *Orthonairovirus* endothelial cells. In that study, reducing the glycoprotein levels with siRNA against the M segment had a greater impact on virus copy number (decrease by about 90%) and release [15]. Moreover, the M segment is the most diverse genome of CCHFV. This diversity may come from how CCHFV uses the vectors and vertebrate hosts in different geographic ranges. Therefore, it is difficult to design general well-functioning inhibitory siRNAs for this segment and many studies found a lower inhibitory effect. Although glycoproteins encoded by the M gene are the most variable portion of the CCHF viruses, some functional domains of the glycoproteins are well conserved [2,31].

In case of CCHFV, the largest of the three segments termed the L segment, encodes an RNA-dependent RNA polymerase (RdRp) that is characterized by several conserved functional regions [2]. Moreover, next to nucleoprotein, L protein drives the processes of transcription and replication that occur in the cytoplasm during the viral replication cycle. Thus, targeting this segment is likely to be an exact strategy. In our study, a remarkable copy number decrease (by 92%) was caused by siL4.

Taken together, our results provide further support for the use of RNA interference-based technique in the development of antiviral drugs against CCHFV infections. Moreover, to our knowledge, this is the first study that used designed siRNAs against CCHFV replication *in vitro* and the first study to provide RNAi solution to all three genomic segments of a nairovirus. Currently, CCHFV constitutes a notable public health concern in our region, with significant geographic expansion in recent decades and growing epidemic potential [21,32,33]. One major limitation of our study is the lack of combinative experiments, however, it well projects future research directions. Combining efficient siRNAs with each other may reveal their potential synergic inhibition effect. Accordingly, the threat of viral infection will increase in the coming years, so any kind of research project aimed at preventing and overcoming a possible infection may be useful. Moreover, we would like to design time-dependent experiments that examine siRNAs efficiency before and after CCHFV infection because they are required for *in vivo* experiments in the future. This study gives novel and important research results for one of WHOs prioritized emerging disease and constitutes a major step for future antiviral development efforts.

## Author Contributions

BSL-4 laboratory processes were made by F.F., M.M., P.H., G.K., B.Z.; preparation of A549 cells was made by M.M., H.P.; siRNA design was made by F.F.; transfection and infection experiments were made by F.F.; cytotoxicity test was made by H.P., F.F.; RT-qPCR experiments were made by F.F.; RT-ddPCR experiments and data analysis were made by L.G., G.K., F.F.; manuscript writing was made by F.F.; supervision: F.J., G.K.

## Acknowledgments

We would like to kindly thank the opportunity to use the QX200 Droplet Digital PCR system (Bio-Rad, CA, USA) and valuable help to the Department of Laboratory Medicine (K.G., L.G.), University of Pécs. We would like to thank Levente Bálint and Jon Eugene Marquette to correct the manuscript’s English language.

## References

1. Ergönül, Ö. Crimean-Congo haemorrhagic fever. Lancet Infect. Dis. 2006, 6, 203–214.

2. Zivcec, M.; Scholte, F.; Spiropoulou, C.; Spengler, J.; Bergeron, É. Molecular insights into Crimean-Congo hemorrhagic fever virus. Viruses 2016, 8, 106.

3. Estrada-Peña, A.; Palomar, A.M.; Santibáñez, P.; Sánchez, N.; Habela, M.A.; Portillo, A.; Romero, L.; Oteo, J.A. Crimean-Congo hemorrhagic fever virus in ticks, Southwestern Europe, 2010. Emerg. Infect. Dis. 2012, 18, 179–80.

4. Keshtkar-Jahromi, M.; Kuhn, J.H.; Christova, I.; Bradfute, S.B.; Jahrling, P.B.; Bavari, S. Crimean-Congo hemorrhagic fever: current and future prospects of vaccines and therapies. Antiviral Res. 2011, 90, 85–92.

5. Fire, A.; Xu, S.; Montgomery, M.K.; Kostas, S.A.; Driver, S.E.; Mello, C.C. Potent and specific genetic interference by double-stranded RNA in Caenorhabditis elegans. Nature 1998, 391, 806–811.

6. Dykxhoorn, D.M.; Lieberman, J. Silencing viral infection. PLoS Med. 2006, 3, e242.

7. Coburn, G.A.; Cullen, B.R. Potent and specific inhibition of human immunodeficiency virus type 1 replication by RNA interference. J. Virol. 2002, 76, 9225–9231.

8. Jacque, J.M.; Triques, K.; Stevenson, M. Modulation of HIV-1 replication by RNA interference. Nature 2002, 418, 435–438.

9. He, M.-L. Inhibition of SARS-associated coronavirus infection and replication by RNA interference. JAMA J. Am. Med. Assoc. 2003, 290, 2665–2666.

10. Shlomai, A.; Shaul, Y. Inhibition of hepatitis B virus expression and replication by RNA interference. Hepatology 2003, 37, 764–770.

11. Kronke, J.; Kittler, R.; Buchholz, F.; Windisch, M.P.; Pietschmann, T.; Bartenschlager, R.; Frese, M. Alternative approaches for efficient inhibition of Hepatitis C virus RNA replication by small interfering RNAs. J. Virol. 2004, 78, 3436–3446.

12. Ge, Q.; McManus, M.T.; Nguyen, T.; Shen, C.H.; Sharp, P.A.; Eisen, H.N.; Chen, J. RNA interference of influenza virus production by directly targeting mRNA for degradation and indirectly inhibiting all viral RNA transcription. Proc. Natl. Acad. Sci. U. S. A. 2003, 100, 2718–2723.

13. Flusin, O.; Vigne, S.; Peyrefitte, C.N.; Bouloy, M.; Crance, J.-M.; Iseni, F. Inhibition of Hazara nairovirus replication by small interfering RNAs and their combination with ribavirin. Virol. J. 2011, 8, 249.

14. Maffioli, C.; Grandgirard, D.; Leib, S.L.; Engler, O. SiRNA inhibits replication of langat virus, a member of the tick-borne encephalitis virus complex in organotypic rat brain slices. PLoS One 2012, 7.

15. Chiang, C.-F.; Albariň, C.G.; Lo, M.K.; Spiropoulou, C.F. Small interfering RNA inhibition of andes virus replication. PLoS One 2014, 9, 99764.

16. Karothia, D.; Dash, P.K.; Parida, M.; Bhagyawant, S.; Kumar, J.S. Inhibition of West Nile virus replication by bifunctional siRNA targeting the NS2A and NS5 conserved region. Curr. Gene Ther. 2018, 18, 180–190.

17. Kim, V.N. Small RNAs: classification, biogenesis, and function. Mol. Cells 2005, 19, 1–15.

18. Tompkins, S.M.; Lo, C.-Y.; Tumpey, T.M.; Epstein, S.L. Protection against lethal influenza virus challenge by RNA interference in vivo. Proc. Natl. Acad. Sci. U. S. A. 2004, 101, 8682–6.

19. McCaffrey, A.P.; Nakai, H.; Pandey, K.; Huang, Z.; Salazar, F.H.; Xu, H.; Wieland, S.F.; Marion, P.L.; Kay, M.A. Inhibition of hepatitis B virus in mice by RNA interference. Nat. Biotechnol. 2003, 21, 639–644.

20. Duh, D.; Nichol, S.T.; Khristova, M.L.; Saksida, A.; Hafner-Bratkovič, I.; Petrovec, M.; Dedushaj, I.; Ahmeti, S.; Avšič-Županc, T. The complete genome sequence of a Crimean-Congo hemorrhagic fever virus isolated from an endemic region in Kosovo. Virol. J. 2008, 5, 7.

21. Földes, F.; Madai, M.; Németh, V.; Zana, B.; Papp, H.; Kemenesi, G.; Bock-Marquette, I.; Horváth, G.; Herczeg, R.; Jakab, F. Serologic survey of the Crimean-Congo haemorrhagic fever virus infection among wild rodents in Hungary. Ticks Tick. Borne. Dis. 2019, 10.

22. Yuan, B.; Latek, R.; Hossbach, M.; Tuschl, T.; Lewitter, F. siRNA Selection Server: an automated siRNA oligonucleotide prediction server. Nucleic Acids Res. 2004, 32, W130–4.

23. Atkinson, B.; Chamberlain, J.; Logue, C.H.; Cook, N.; Bruce, C.; Dowall, S.D.; Hewson, R. Development of a real-time RT-PCR assay for the detection of Crimean-Congo hemorrhagic fever virus. Vector-Borne Zoonotic Dis. 2012, 12, 786–793.

24. R Core Team R: a language and environment for statistical computing 2019.

25. Wickham, H. Getting started with qplot BT-ggplot2: elegant graphics for data analysis. In ggplot2; Wickham, H., Ed.; Springer New York, 2009; pp. 9–26 ISBN 978-0-387-98141-3.

26. Levin, A.; Kutznetova, L.; Kahana, R.; Rubinstein-Guini, M.; Stram, Y. Highly effective inhibition of Akabane virus replication by siRNA genes. Virus Res. 2006, 120, 121–127.

27. Flusin, O.; Vigne, S.; Peyrefitte, C.N.; Bouloy, M.; Crance, J.M.; Iseni, F. Inhibition of Hazara nairovirus replication by small interfering RNAs and their combination with ribavirin. Virol. J. 2011, 8.

28. Das, A.T.; Brummelkamp, T.R.; Westerhout, E.M.; Vink, M.; Madiredjo, M.; Bernards, R.; Berkhout, B. Human Immunodeficiency Virus Type 1 Escapes from RNA Interference-Mediated Inhibition. J. Virol. 2004, 78, 2601–2605.

29. Garcia, S.; Billecocq, A.; Crance, J.-M.; Munderloh, U.; Garin, D.; Bouloy, M. Nairovirus RNA sequences expressed by a Semliki Forest virus replicon induce RNA interference in tick cells. J. Virol. 2005, 79, 8942–8947.

30. Billecocq, A.; Vazeille-Falcoz, M.; Rodhain, F.; Bouloy, M. Pathogen-specific resistance to Rift Valley fever virus infection is induced in mosquito cells by expression of the recombinant nucleoprotein but not NSs non-structural protein sequences. J. Gen. Virol. 2000, 81, 2161–2166.

31. Erickson, B.R.; Deyde, V.; Sanchez, A.J.; Vincent, M.J.; Nichol, S.T. N-linked glycosylation of Gn (but not Gc) is important for Crimean Congo hemorrhagic fever virus glycoprotein localization and transport. Virology 2007, 361, 348–355.

32. Hornok, S.; Horváth, G. First report of adult Hyalomma marginatum rufipes (vector of Crimean-Congo haemorrhagic fever virus) on cattle under a continental climate in Hungary. Parasit. Vectors 2012, 5, 170.

33. Németh, V.; Oldal, M.; Egyed, L.; Gyuranecz, M.; Erdélyi, K.; Kvell, K.; Kalvatchev, N.; Zeller, H.; Bányai, K.; Jakab, F. Serologic evidence of Crimean-Congo hemorrhagic fever virus infection in Hungary. Vector-Borne Zoonotic Dis. 2013, 13, 270–272.

